# Mapping Niche-specific Two-Component System Requirements in Uropathogenic *Escherichia coli*

**DOI:** 10.1101/2023.05.23.541942

**Authors:** John R. Brannon, Seth A. Reasoner, Tomas A. Bermudez, Taryn L. Dunigan, Michelle A. Wiebe, Connor J. Beebout, Tamia Ross, Adebisi Bamidele, Maria Hadjifrangiskou

## Abstract

Sensory systems allow pathogens to differentiate between different niches and respond to stimuli within them. A major mechanism through which bacteria sense and respond to stimuli in their surroundings is two-component systems (TCSs). TCSs allow for the detection of multiple stimuli to lead to a highly controlled and rapid change in gene expression. Here, we provide a comprehensive list of TCSs important for the pathogenesis of uropathogenic *Escherichia coli* (UPEC). UPEC accounts for >75% of urinary tract infections (UTIs) worldwide. UTIs are most prevalent among people assigned female at birth, with the vagina becoming colonized by UPEC in addition to the gut and the bladder. In the bladder, adherence to the urothelium triggers *E. coli* invasion of bladder cells and an intracellular pathogenic cascade. Intracellular *E. coli* are safely hidden from host neutrophils, competition from the microbiota, and antibiotics that kill extracellular *E. coli.* To survive in these intimately connected, yet physiologically diverse niches *E. coli* must rapidly coordinate metabolic and virulence systems in response to the distinct stimuli encountered in each environment. We hypothesized that specific TCSs allow UPEC to sense these diverse environments encountered during infection with built-in redundant safeguards. Here, we created a library of isogenic TCS deletion mutants that we leveraged to map distinct TCS contributions to infection. We identify – for the first time – a comprehensive panel of UPEC TCSs that are critical for infection of the genitourinary tract and report that the TCSs mediating colonization of the bladder, kidneys, or vagina are distinct.

**IMPORTANCE:** While two-component system (TCS) signaling has been investigated at depth in model strains of *E. coli*, there have been no studies to elucidate – at a systems level – which TCSs are important during infection by pathogenic *Escherichia coli*. Here, we report the generation of a markerless TCS deletion library in a uropathogenic *E. coli* (UPEC) isolate that can be leveraged for dissecting the role of TCS signaling in different aspects of pathogenesis. We use this library to demonstrate, for the first time in UPEC, that niche-specific colonization is guided by distinct TCS groups.

## INTRODUCTION

Whether to newly colonized niches or to changing conditions, bacteria efficiently adapt to environmental changes by rapidly changing gene expression (1, 2). A major mechanism through which bacteria interpret environmental changes into specific changes in their gene expression is two-component signal transduction systems (TCSs). TCSs are usually comprised of two parts: a sensor histidine kinase (HK) and a cognate response regulator (RR). Typically, signal interception results in HK autophosphorylation at a conserved histidine residue and subsequent phosphotransfer to a conserved aspartate on the RR. The most widely observed outcome of RR phosphorylation is increased DNA binding affinity of the phosphorylated RR for its target promoters, consequently altering target gene expression.

Compartmentalized, TCSs serve as bacterial logic gates that process sensory input with the net output result often being a change in gene expression that ultimately changes one or multiple bacterial phenotypes. However, multiple studies show that – like in mammalian signaling systems – TCSs are dynamic and branch to incorporate multiple stimuli, interact outside the boundaries of cognate partners, or be part of phosphorelays that allow for a refined, beneficial orchestration of molecular systems (3). While these phenomena are well-characterized *in vitro*, either in biochemical protein-protein interactions or in the test tube, few studies have investigated how each TCS of a given pathogen contributes to host colonization.

Uropathogenic *E. coli* (UPEC) is the causative agent of ∼80% of the 150 million urinary tract infections (UTIs) that occur annually (4–6). UPEC’s ability to survive within several different environments contributes to its successful prevalence. UPEC can be transmitted amongst individuals through the fecal-oral route and sexual contact (4). Within the intestinal tract, UPEC colonizes the human gut alongside commensals for extended periods of time, in contrast to diarrheagenic *E. coli* pathotypes (6–8). However, unlike commensals, the genetically diverse UPEC strains are equipped with fitness determinants that allow them to expand beyond the gut (8–11). Exit from the gut, is followed by urethral accession to the bladder causing cystitis. During bladder infection, UPEC dynamically reach the kidney and in some cases can cause pyelonephritis, from where bacteria can traverse to the bloodstream, leading to bacteremia (12–14). In people assigned female at birth, who are disproportionally impacted by UTIs, UTI pathogenesis additionally encompasses the vagina (15, 16). Within these different host niches, UPEC is found in extra- and intracellular compartments exposed accordingly to different stresses and metabolite inventories. In the bladder and vaginal lumens, UPEC exist planktonically or associated with the urothelial or vaginal cells. Likewise in the kidney, UPEC associates with the host cell membrane (17). In the host intracellular environments, UPEC form three different intracellular bacterial reservoirs: metabolically active, biofilm-like, intracellular bacterial communities (IBCs) in the superficial umbrella cells of the bladder; quiescent intracellular reservoirs inside transitional bladder cells and vaginal intracellular communities within vaginal epithelial cells (15, 18).

While the field has developed an in-depth understanding of the molecular systems contributing to UPEC’s plasticity, such as flagella curli, and type 1 and P pili, little is known about which TCSs are coordinately used during infection. In this work, we tested the hypothesis that the different unique niches UPEC encounters necessitate the use of distinct TCSs. To test this hypothesis, we constructed an isogenic deletion mutant library of all the TCSs encoded by the prototypical cystitis isolate UTI89 (**Table 1**) that is extensively used in the field to study UTI pathogenesis (19). As this is the first time a comprehensive TCS deletion library has been constructed for study in a UPEC isolate, an in-depth *in vitro* analysis of how each TCS deletion strain associates with bladder cells is was first performed. Next, we leveraged our well-established UTI mouse models to determine the pathogenicity of each TCS deletion mutant in the bladder, kidneys, and vagina. We report that *in vitro*, none of the TCSs deletion mutants substantially influences adherence or invasion of urothelial cells, indicating that this critical step in infection establishment is under redundant control. Our *in vivo* data uncover – for the first time – the inventory of TCSs needed for bacterial expansion following the adherence and invasion steps in the bladder. These TCSs are distinct from the TCSs that we found to be critical for kidney or vaginal colonization following bladder infection. Collectively, our results demonstrate niche-specific requirements for TCSs during the different stages of UTIs and lay the groundwork for delineating mechanistic details associated with each network during UPEC infection.

**Table 1:** Information on HK – RR pairs and orphan proteins in UPEC strain UTI89. All information was compiled using P2CS and MiST3.0 databases.

| **HK** | **Loci** | **RR** | **Loci** | **Ligand or stimulating conditions** | **Ref.** |
| --- | --- | --- | --- | --- | --- |
| ArcB | C3646 | ArcA | C5174 | Anaerobic growth | (51) |
| AtoS | C2501 | AtoC | C2502 | Acetoacetate | (52) |
| BaeS | C2353 | BaeR | C2354 | Envelope stress, indole, and flavonoids | (53) |
| BarA | C3156 | UvrY | C2115 | Acetate and short-chain fatty acids | (54) |
| BtsS | C2398 | BtsR | C2397 | Pyruvate | (55) |
| / | C4639 | / | C4638 | Unknown | / |
| / | C4932 | / | C4931 | Unknown | / |
| / | C4937 | / | / | Unknown | / |
| CpxA | C4495 | CpxR | C4496 | Envelope stress | (56) |
| CreC | C5170 | CreB | C5171 | During glucose fermentation | (57) |
| CusS | C0570 | CusR | C0571 | Copper and silver | (58) |
| DcuS | C4719 | DcuR | C4718 | Presence of C4-dicarboxylates like malate | (59, 60) |
| DpiB | C0624 | DpiA | C0624 | Citrate | (61) |
| EnvZ | C3904 | OmpR | C3905 | Changes in osmolarity | (62) |
| EvgS | C2702 | EvgA | C2701 | Alkali metals and low pH | (63) |
| GlnL | C4458 | GlnG | C4457 | Nitrogen deprivation | (64) |
| KdpD | C0699 | KdpE | C0698 | Potassium limitation | (65) |
| NarQ | C2795 | / | / | Nitrate and nitrite | (66) |
| NarX | C1418 | NarL | C1417 | Nitrate and nitrite | (66, 67) |
| PhoQ | C1258 | PhoP | C1259 | Low magnesium and calcium and membrane stress | (68, 69). |
| PhoR | C0421 | PhoB | C0420 | Inorganic phosphate | (70) |
| PmrB | C4706 | PmrA | C4707 | Ferric iron and membrane stress | (71) |
| QseC | C3451 | QseB | C3450 | Unknown | / |
| QseE | C2876 | QseF | C2873 | Adrenergic receptor | (72) |
| RcsCD | C2500,C2498 | RcsB | C2499 | Osmotic shock | (73) |
| RstB | C1796 | RstA | C1797 | Unknown | / |
| TorS | C1056 | TorR | C1059 | Trimethylamine-N-oxide | (74) |
| UhpB | C4224 | UhpA | C4225 | Glucose-6-phosphate | (75) |
| YedV | C2167 | YedW | C2168 | Hydrogen peroxide responsive | (76) |
| YpdA | C2712 | YpdB | C2713 | High pyruvate concentrations | (55) |
| ZraS | C3816 | ZraR | C3815 | Zinc | (77) |
|  |  | NarP | C2471 | Nitrate and nitrite | (66) |
|  |  | RssB | C1431 | Glucose, phosphate, and nitrogen deprivation | (78) |

## METHODS

### Ethics Statement

Ethics approval for mouse experiments was granted by the VUMC Institutional Animal Care and Use Committee (IACUC) (protocol numbers M1800101-01). Experiments were performed in accordance with the guidelines of the National Institute of Health and IACUC at VUMC.

### Growth of Eukaryotic Cells and Bacterial strains

A complete list of *E. coli* strains and their characteristics used for this study are listed in Table S1. From freezer stocks, strains were grown in 5 mL of lysogeny broth (LB) at 37 °C with shaking. For bacteriological assays, LB culture tubes were incubated overnight. For inoculation of 5637 (ATCC HTB-9) cells and C3H/HeN mice, bacterial cultures were first seeded from a freezer stock and incubated for 4 h at 37 °C, followed by two sequential, ∼24h sub-cultures (1:1000 dilution) in 10 mL, static LB media at 37 °C in-order to induce type 1 pili (19). Preceding inoculation of immortalized cells and mice, strains were normalized in 1x PBS. The immortalized bladder epithelial cell line 5637 (ATCC HTB-9) was grown statically at 37 °C in RPMI 1640 (Life Technologies Co., Grand Island, NY) media supplemented with 10% FBS under 5% CO_2_.

### Construction of UPEC TCS Deletion Library

A complete list of plasmids and primers can be found in Tables S2 and S3, respectively. For our study, we used the genetically tractable, cystitis strain UTI89 and using the prokaryotic TCS databases, P2CS and MiST, identified a comprehensive list of TCSs in UTI89 for deletion (20, 21). Targeted TCS deletion mutants for the isogenic deletion library were constructed using the λ Red recombinase system and gene deletion was confirmed by PCR with test primers (22, 23).

### Bacterial Growth Curves

Strains were grown overnight in LB broth and sub-cultured to a starting OD_600_ of 0.05 in the specified broth in a 96-well plate. Plates were incubated at 37°C with shaking for 8 h and OD_600_ readings taken every 15 min. Growth data was fit to the Weibull growth model to determine specific growth rate (24).

### Gentamicin-Based Protection Assays

Experiments were performed as described elsewhere (25). 5637 (ATCC HTB-9) bladder epithelial cells were grown to at least 90% confluency in 24-well plates. Prior to inoculation with *E. coli*, new RPMI 1640 media supplemented with 10% was added to the cells. To achieve an approximate MOI of 5, 5637 cell density was enumerated to determine the amount of *E. coli* suspension to be used per well. Strain inoculum was added to three sets of triplicate wells and centrifuged for 5 min at 600×*g* to facilitate uniform contact between bladder and bacteria cells. Following, plates were incubated at 5% CO_2_ and 37 °C for 2 h. For cell lysis, a final concentration of 0.1% Triton X-100 was used. Bacterial burden was enumerated by serial dilution and spot platting onto LB plates. One set of wells was lysed to determine the total number of bacteria within the well. The other two sets were washed with 0.5 mL of PBS three times. The *E. coli* adhering to bladder cells was enumerated in a set of wells that was immediately lysed after the washes. The final set of wells were gently washed with PBS with 100 μg/mL of gentamicin (Life Technologies Co., Grand Island, NY) for 2h; afterwards, wells were washed two more times with 1 mL of PBS to enumerate the intracellular *E. coli*. The percent of *E. coli* adherence and invasion were calculated as a percentage of the total number of bacteria.

### Mouse UTI Model

Mouse infectious were performed as previously described (26, 27). *E. coli* strains were incubated at 37 °C, initially in a 5 mL LB culture tube with shaking for 4 h and followed by two sequential sub-cultures at 1:1,000 into 10 mL of fresh LB and grown statically for 24 h. C3H/HeN female mice at 7-to 8-weeks-old were transuretherally inoculated with 50 µL an *E. coli* suspension of 10^7^ CFUs in PBS and mice were humanly euthanized at 6 h, 24 h, or 28 days post-infection (h/dpi). After euthanasia, organs were removed and homogenized in PBS for CFU enumeration. For quantification of intracellular bacteria, bladder tissue was incubated in 100 μg/mL of gentamicin for 2 h, washed with PBS, and homogenized in PBS with 0.1% Triton X-100.

### Visualization and Enumeration of intracellular bacterial communities

IBC enumeration was performed as described previously using bacterial strains transformed with pCOM::GFP plasmid (13). Mice were euthanized 6 hpi and the bladders were removed with aseptic technique. Mouse bladders were stretched, pinned, and fixed with 3.4% paraformaldehyde overnight at 4 °C. Bladders were washed twice in 1X PBS, permeabilized with 0.1% Triton X-100 for 15 min, followed by a 1XPBS wash. Bladders were stained at room temperature with Alexa Fluor 568 Phalloidin (ThermoFisher) and mounted on to slides with ProLong Diamond Antifade (ThermoFisher). IBCs were manually counted via fluorescence microscopy on a LSM 710 confocal laser scanning microscope (Zeiss).

### Statistical Analysis

Statistical analyses were performed in GraphPad Prism software, using the most appropriate test for each analysis. Experiments were performed in accordance with the standard convention, incorporating at least three biological replicates. Statistical tests used for analysis are two-tailed. Additionally, for mouse infections, power analyses were performed to determine seven subjects per group are needed to achieve a power level of 90% for detecting a 25% difference in the CFU means with an in-group standard deviation of 20%. Additional experimental details of group size, statistical test, error bars, and probability value for each statistically evaluated experiment are specified in the corresponding figures, legends, and text.

## RESULTS

### Construction of a UPEC TCS deletion library

Using the P2CS and MiST3.0 databases, we compiled a list of TCS within the UPEC strain UTI89. In UTI89, we found 32 RRs and HKs, including four hybrid HKs (**Table 1**). Typically, classical TCSs components are encoded together in an operon; however, in some cases TCS component genes are encoded at distinct loci in the chromosome or comprise more complex multi-branch systems or phosphorelay systems, such as RcsCDB. Finally, certain strains, including UTI89, may encode “orphan” TCS components with no known interaction partners. Within UTI89, we identified 25 orthodox pairs and 14 orphans, including one without a known partner. In order to construct a comprehensive TCSs deletion library in UTI89 accounting for gene separation, we generated 38 different isogenic deletion mutants. Orthodox TCSs were deleted in pairs, and orphans were deleted separately. For analysis, the *rcsDB* gene cluster that codes for the phosphorelay system RcsDBC, were deleted as a unit together. The unorthodox CheA-CheY chemosensory system, which has been heavily investigated in the context of chemotaxis (28, 29), was omitted from this study. All resulting mutant strains are marker-less and validated by PCR. To our knowledge, this is the first time a TCS deletion library is constructed in a UPEC isolate.

### Deletion of a singular TCS pairs does not impair adherence or invasion of urothelial cells

Prior to evaluating the contribution of each TCS to UPEC pathogenesis, we sought to determine whether deletion of any TCS negatively affects growth, adherence or invasion. To assess strains for their contribution to growth *in vitro*, we performed growth curves in nutrient limited N-minimal growth media or in the nutrient-rich lysogeny broth (LB). Growth curves were fit to the Weibull growth model to compare specific growth rates. The specific growth rate of UTI89 was 0.1962 ±0.0542 h^-1^ for N-minimal (**Fig. 1A**) and 0.1926 ±0.0222 h^-1^ for LB media (**Fig. 1B**) (mean ±95% CI) with no statistically significant difference compared to any of the TCS mutant strains tested). These results indicate that – under the conditions tested – not a single TCS is critical to planktonic growth of UPEC strain UTI89. We thus proceeded to evaluate the TCS deletion library on specific phenotypes associated with the initial stages of infection.

**Figure 1:**
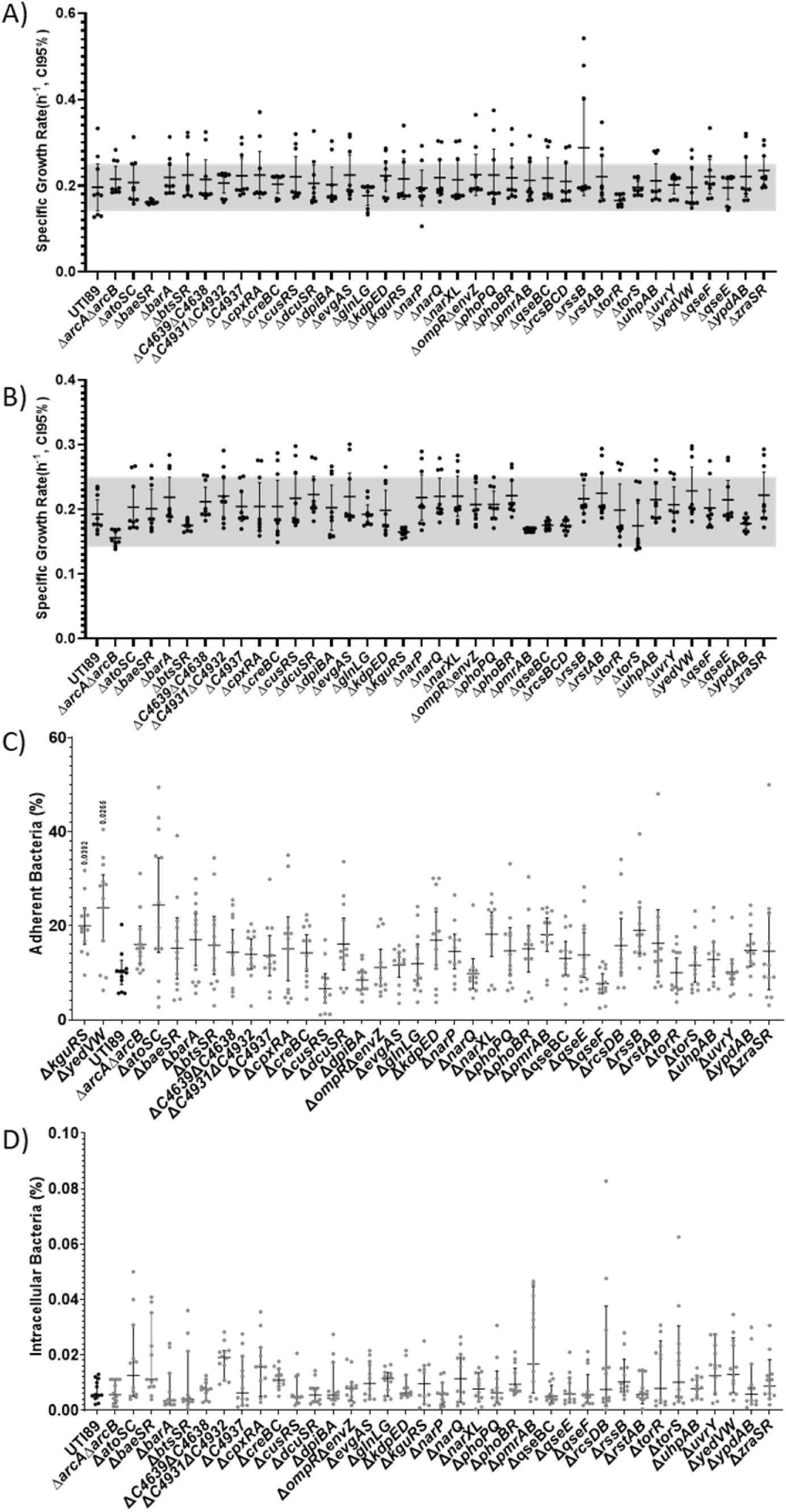
*In vitro* properties of UPEC TCS deletion mutants. A) Graph depicts the specific growth rate of each TCS mutant compared to the isogenic parent UTI89, during growth in (**A**) N-minimal or (**B**) LB media with shaking, at 37°C. Growth curves were fit to the Weibull growth model to determine specific growth rate. (**C**) Adherent and (**D**) intracellular bacterial titers for wild-type UTI89 and each of the isogenic TCS deletion strains. Experiments were performed using an MOI of 5 on the immortalized urothelial cell line 5637. The percent of *E. coli* adherence and invasion were calculated as a percentage of the total number of bacteria with a well at the 2 h endpoint. Each dot represents a biological replicate. A non-parametric Kruskal–Wallis with two-sided Dunn’s post-hoc test was performed for statistical analysis.

UPEC infection begins with adherence and invasion of urothelial cells. This process is governed by type 1 pili (*fim*), which are adhesive fibers assembled by the chaperone-usher pathway. A defect in adherence excludes invasion of the urothelium and constitutes a bottlenecking event. To determine if any of the UPEC TCS mutants display an adherence or invasion defect, we leveraged a well-established tissue culture model (25) using the 5637 immortalized urothelial cell line. These assays revealed that Δ*yedVW* and Δ*kguRS* mutants had adherence levels higher than the parental UTI89 strain (**Fig. 1C**). Nonetheless, enumeration of internalized bacteria did not identify statistically significant differences among strains (**Fig. 1D**). These data again indicate that no single TCS deletion impairs the initial steps in UPEC pathogenesis that typically acts as a bottleneck for studying downstream infection stages. Knowing that none of the TCS deletion strains has an adherence or invasion defect, we went on to test the entire deletion library in a murine UTI model.

### Distinct TCSs contribute to colonization of genitourinary tract niches

We next sought to evaluate the contribution of each TCS mutant in a UTI mouse model. In this model, UPEC adheres and invades urothelial cells, ascends to the kidney, and migrates to the vagina and gut in a dynamic fashion (26, 30, 31). Acute infection hallmarks include the formation of IBCs at 6 hpi and bacteriuria that persists over time in ∼50% of the infected mice (27). Another hallmark of UTI that is captured in this murine model, is the formation of asymptomatic reservoirs in the vagina (30).

In this experiment, we asked how each TCS mutant colonizes three niches: the bladder, kidneys, and vagina (30). Cohorts of 6-8 week old female C3H/HeN mice were transuretherally inoculated with the wild-type parent or each of the isogenic TCS mutants. Mice were euthanized 24 hpi and bacterial titers in the bladder, kidneys, and vagina were enumerated for each infected mouse. These experiments revealed 12 different TCS deletion strains with altered bacterial titers (**Fig. 2**). Consistent with the hypothesis that distinct TCS are needed in unique sub-niches in the genitourinary tract, we observed mutants with niche-specific defects.

**Figure 2:**
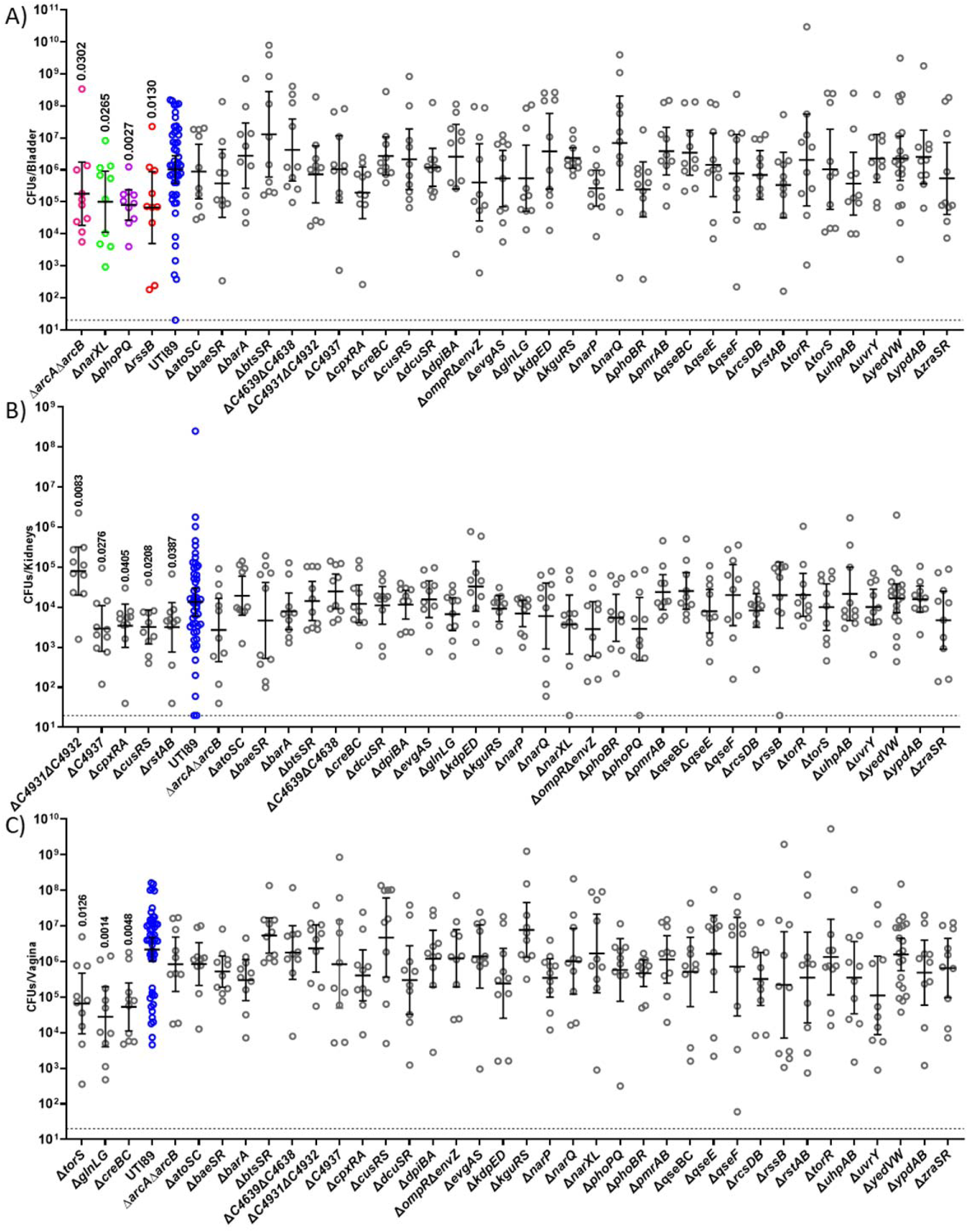
Niche-specific contribution of TCSs during 24h infection: Mice infected with UTI89 (Blue) and isogeneic TCS deletion strains were euthanized 24 hpi for bacterial enumeration of titer within the (**A**) bladder, (**B**) kidneys, and (**C**) vagina. Each dot represents organ titers from a different mouse. A non-parametric Kruskal–Wallis with two-sided uncorrected Dunn’s post-hoc test was performed for statistical analysis.

In the bladder, the Δ*arcA*Δ*arcB*, Δ*narXL*, Δ*phoPQ* and Δ*rssB* showed decreased titers compared to the parent strain at 24 hpi (**Fig. 2A**), but were able to colonize the kidney and transit to the vagina similar to the wild-type strain (**Fig. 2B-C**). Conversely, ΔC4931ΔC4932, ΔC4937, Δ*cpxRA*, Δ*cusRS*, Δ*rstAB* exhibited altered titers in the kidney (**Fig. 2B**), while Δ*torS*, Δ*creBC*, Δ*glnLG* exhibited significant colonization defects in the vagina (**Fig. 2C**) at 24 hpi. These different TCS make apparent contributions to the respective niches within the acute mouse model. Moreso, these results indicate that while these TCSs contribute to survival within a niche, they are dispensable for survival of the other two niches.

### Mutants defective for bladder colonization are associated with energy metabolism

Our lab has previously elucidated that aerobic respiration is critical for intracellular replication of UPEC (32). The current analyses uncover 4 TCS mutants, Δ*arcA*Δ*arcB*, Δ*narXL*, Δ*phoPQ*, and Δ*rssB* RR that display defects in bladder colonization at 24 hpi. ArcA/ArcB, NarX/NarL and RssB belong to regulatory networks associated with respiration (33, 34), while PhoP/PhoQ is implicated in stress response and energy metabolism (35, 36) (**Fig. 2A**). We therefore focused on these regulators to further dissect the stage at which they become important during infection and to evaluate how their deletion impacts long-term persistence of UPEC in the urinary tract.

During UTI, UPEC become internalized by urothelial cells, in which they replicate into biofilm-like communities by consuming oxygen primarily via the quinol oxidase cytochrome bd (37). Previous work demonstrated that the activity of the ArcB HK is influenced by the quinol oxidation state and that both RssB and ArcA influence the abundance of the sigma factor σ^38^ (RpoS) that in turn influences the expression of biofilm components in *E. coli* (33, 38–41). To determine whether each bladder-defective TCS deletion mutant has the ability to form IBCs during acute infection, we evaluated intracellular bacterial titers and IBC formation at 6 hpi. To enumerate the extracellular and intracellular bladder populations, mice were euthanized at 6 hpi, and bladders were removed, bisected, and gentamicin-treated to eliminate extracellular bacteria and enable enumeration of the intracellular bacterial levels. These analyses revealed that all four mutants, Δ*arcA*Δ*arcB*, Δ*narXL*, Δ*phoPQ*, and Δ*rssB*, are retained in the bladder lumen, with titers similar to the wild-type parent (**Fig. 3A**); in contrast to the extracellular fractions, the intracellular titers of the mutants were statistically significantly different than the parental strain titers. The mutants Δ*arcA*Δ*arcB*, Δ*narXL*, and Δ*phoPQ* had high intracellular titers, whereas the Δ*rssB* mutant has drastically lower titers than the parental UTI89 strain (**Fig. 3B**) that are reminiscent of a mutant that lacks cytochrome bd (37). Based upon the 6 hpi bacterial titer analysis, we sought to determine whether the *rssB* or *arcAB* deletion resulted in altered IBC morphology or numbers. We performed microscopy on these selected strains at 6 hpi. We did not observe any apparent difference in IBC morphology or size (**Fig. S1**). We noted an abundance of IBCs in the Δ*arcA*Δ*arcB* strain (**Fig. 3C**), which is consistent with the observed increase in intracellular bacterial titers. Conversely, there was a statistically significant lower number of IBCs in the Δ*rssB* infected bladders compared to the UTI89 strain (**Fig. 3C**), in agreement with the significantly lower numbers of intracellular numbers observed for the Δ*rssB* mutant. This experiment demonstrates that during the intracellular stage of infection ArcA/ArcB and RssB likely contribute to different aspects of the intracellular pathogenic cascade.

**Figure 3:**
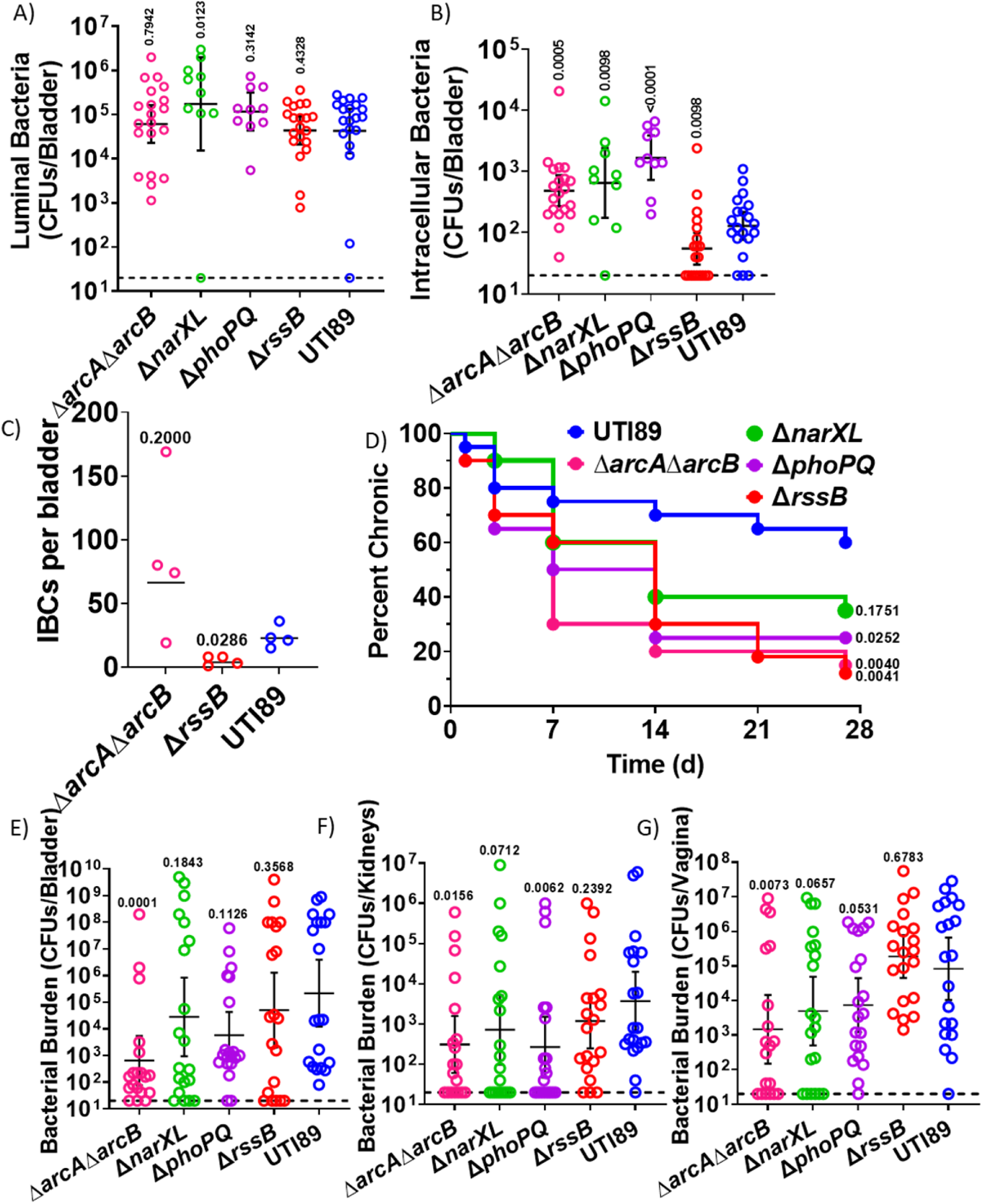
TCSs contribute to distinct stages of extracellular and intracellular pathogenesis. **A-B**) Graphs depict the (**A**) luminal and (**B**) intracellular bacterial titers from mouse bladders assessed at 6 hpi. (**C**) Graph depicts the number of IBCs enumerated in each infected bladder at 6 hpi. Enumeration of IBCs was performed using confocal microscopy on randomly selected bladders. Statistical analysis was performed with Mann Whitney U test. **D-G**) Strains with aberrant acute infection titers were assessed in a long-term 28-day UTI model. **D**) Graph depicts time-to-resolution curves as defined as urine bacterial titers dropping below 10^4^ CFUs/mL. Time to event was modeled with Kaplan-Meier method with non-parametric Mantel-Cox test for statistical comparison of TCS deletion mutants to UTI89. Post 28-day urine analysis, bacterial titers were enumerated in the mouse (**E**) bladder, (**F**) kidneys, and (**F**) vagina. Each symbol is a mouse. Statistical analysis was performed with Mann Whitney U test.

Following acute infection, C3H/HeN mice may develop chronic cystitis or the infections may resolve as indicated by bacterial urine titers dropping below 10^4^ CFUs/mL. Chronic or resolved infection depends upon the bacteria clearing an early host-pathogen checkpoint within the first, 24 hours of acute infection (42, 43). We followed the urine titers of mice infected with either UTI89, Δ*arcA*Δ*arcB*, Δ*narXL*, or Δ*phoPQ* for 28 dpi and measured bacterial organ titers on day 28. Longitudinal urinalysis showed that despite harboring higher numbers of intracellular bacteria at the 6 h time-point, mice infected with Δ*arcA*Δ*arcB*, had a shorter time to resolution event, compared to wild-type UTI89 (**Fig. 3D**). Similarly, urine analysis showed that the Δ*phoPQ*, and Δ*rssB* infected mouse cohorts had a shortened time to resolution compared to those infected with UTI89 (**Fig. 3D**). At the 28-dpi endpoint, the Δ*arcA*Δ*arcB* mutant was the only strain that was statistically significant in lower kidneys, bladder, and vagina titers than the parental UTI89 strain (**Fig. 3E-G)**. The Δ*phoPQ* strain was significantly lower in the kidneys (**Fig. 3F**).

On a broad scale, these data indicate that TCSs make niche and time specific contributions to dynamic UPEC pathogenesis. On a finer scale, these data begin to indicate that UPEC may need to alternate between aerobic and anaerobic respiration states during different stages of infection. Collectively, our study elucidates the niche-specific TCS requirements of UPEC infection (**Fig. 4**); knowledge that we believe will seed future research on better understanding how UPEC regulate virulence and fitness determinants in a dynamic manner.

**Figure 4:**
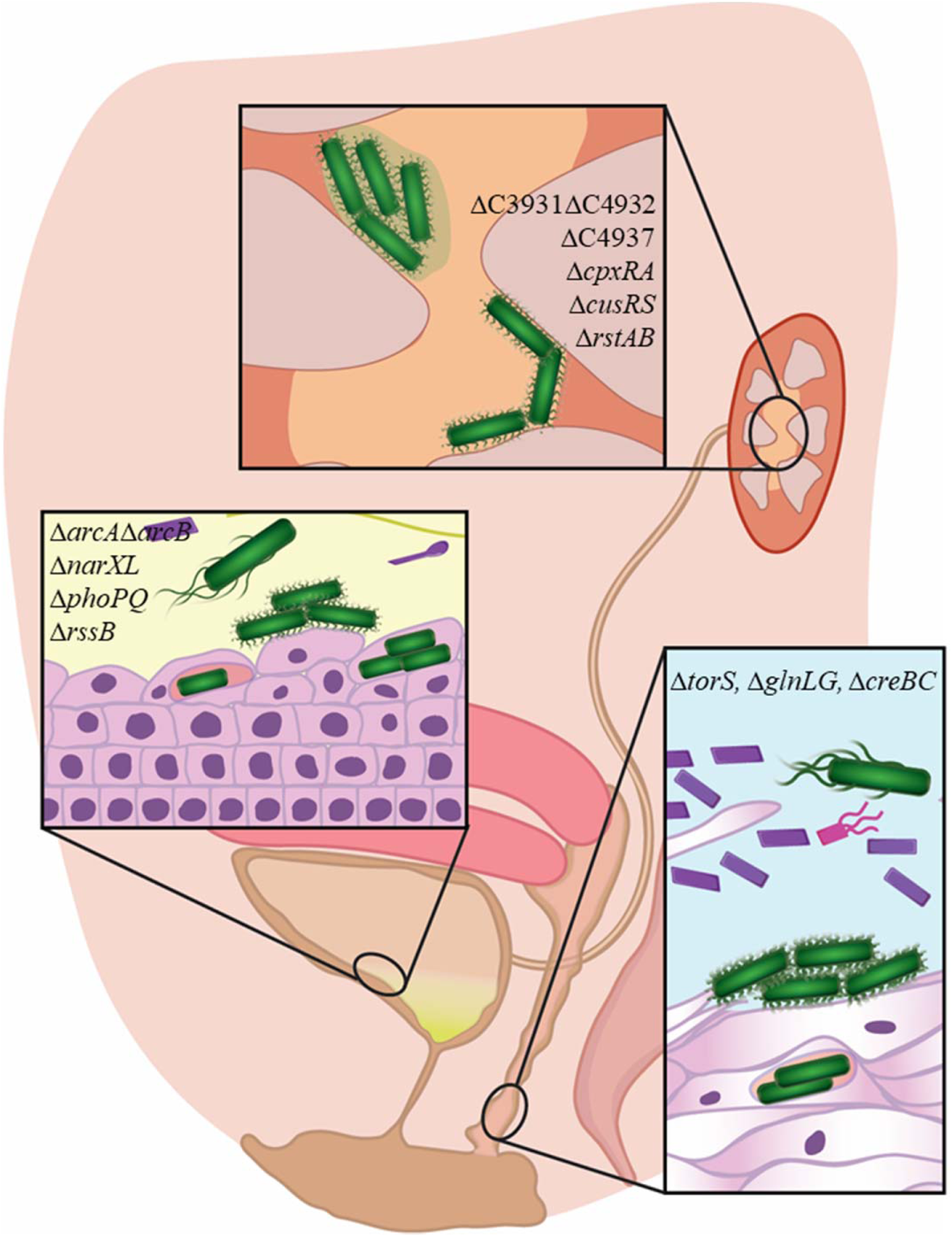
Summary of the different TCS and their relevant niches. Overall, our data support redundant regulatory networks within TCSs. Among the many TCSs that UPEC encodes, 12 TCSs make critical, niche-specific contributions to UPEC pathogenesis.

## DISCUSSION

A key component to the success of UPEC as a pathogen of the urinary tract and a commensal of the gut and vagina, is its ability to thrive within a variety of different niches and in different states. While studies have thus far extensively focused on virulence factors and the UPEC pathogenic cascade itself, very few reports have investigated regulatory pathways associated with UPEC pathogenic potential. TCSs are a major regulatory mechanism that allow bacteria to switch between molecular tools in response to varying stimuli in their current environment. No studies have been executed to define how UPEC uses TCSs during infection. Here, we generated a deletion library of all the TCSs in the prototypical UPEC strain UTI89 as a tool to probe mechanisms of pathogenesis and persistence. We hope the future use of this library in the field enhances research in this area.

Our study highlights two critical aspects of UPEC TCSs: 1) they’re utilized under specific conditions and 2) likely form complex networks. While, there were no major differences between deletion strains during *in vitro* growth; clear molecular tissue tropisms were observed with *in vivo* experiments for a subset of TCSs. While different TCSs have been expansively studied at the molecular level in *E. coli* and context of gut colonization, only a few have been studied in the context of UPEC pathogenesis. Previous studies have connected a few TCSs to UPEC pathogenesis with a targeted individual approach. For instance, deletion of the *ompR* RR in NU149, which impacts *fim* expression, resulted in approximately 2-log reduction in bladder and kidney titers in BACLB/c mice (44). Deletion of the CpxAR system diminished UTI89 fitness in the bladder after 3 dpi in CBA/J mice (45). The BarA-UvrY system in UPEC CFT073 was reduced in bladder and kidney titers 3 dpi which was attributed to LPS and hemolysin production (46). Deletion of *qseC* alone was documented to lead to UTI89’s attenuation in the mouse bladder (47). These prior studies investigated single component deletions of either the RR or the HK, which can unmask non-partner interactions across TCSs. For this study, we generated a comprehensive TCS deletion library of each TCS pair with the exception or orphan components like RssB, or systems in which the deletion of both TCS partners proved difficult to obtain (*barA*/*uvrY* and *torR*/*torS*). This constructed inventory of TCS deletion mutants now allows for an expansive assessment of the TCSs and their downstream regulons within a variety of settings. In an acute UTI model, we found 12 different systems contributing to infection in either the bladder, kidney, or vaginal niches, focusing at 24 hpi (**Fig. 2**). The library can be leveraged to evaluate the fitness of the TCS mutants at different stages of infection. Moreover, future studies can determine how the fitness potential of each TCS mutant changes if they are inoculated directly in the asymptomatic niches (vagina or gut) instead of transurethrally instilled in the bladder. Will different TCSs become important for exiting the asymptomatic reservoirs and ascending the urethra to the bladder? We think so.

We acknowledge that in this study, we provide a broad view of a single cystitis isolate: the prototypical strain UTI89. UPEC genomic heterogeneity is extensive, and we posit that changes in genomic content may influence TCS use in a strain-specific fashion. Yet, we support that this study is significant as it provides the first comprehensive overview of TCSs to UPEC pathogenesis and lays the foundation for future in-depth investigations in other UPEC isolates and a comparison tool that is more representative than the model laboratory K-12 strains.

In the current study, and based on the expertise of our group, we selected to focus more closely with those TCS mutants that displayed a significant colonization defect in the bladder: D*arcA*D*arcB*, D*narXL*, D*phoPQ* and D*rssB.* (**Fig. 2A, 4**). Three of these mutants, D*arcA*D*arcB*, D*rssB,* and D*narXL* are convergent on the regulation of respiration, a critical aspect of UPEC pathogenesis. Under microaerobic conditions ArcA is connected to the up-regulation of *cydAB* and down-regulation of *cyoABCDE* (48, 49). Recently, *cydAB,* which encodes the cytochrome *bd* oxidase, was found critical for proper IBC expansion (32). The expression of *cydAB* and *arcA* are also regulated by the global one component regulator FNR, which is active at low oxygen levels (49, 50). Along with FNR, the HK NarX, which helps to discern between nitrate and nitrite, phosphorylates NarL which in turn regulates *narG* expression, which is a nitrate reductase. The cytochrome *bd* oxidase and nitrate reductase renew the ubiuinone:ubiquinol pool though be it under different conditions (34). The HK, ArcB controls the phosphorylated state of the RRs ArcA and RssB that co-regulate the balance in RpoS abundance in response to general stress like carbon starvation (33). While our results indicate that the TCS important to cystitis seem to converge on respiration the details to their direct or indirect interactions down or upstream of one another remain to be explored. Further study may reveal details of interconnectivity of these TCSs involved in an energetics balancing act.

In sum, two-component systems are critical systems that mediate a pathogen’s ability to adapt behavior in response to external stressors. TCSs have global impacts on metabolism and virulence factors or targeted towards a narrow regulon. This work provides a comprehensive tool to dissect UPEC pathogenesis from a regulation standpoint and highlight that TCSs are important for UPEC pathogenesis in a niche specific manner.

## ACKNOWLEDGEMENTS

This work was supported by National Institutes of Health (NIH) grants T32GM007569 (JRB), R01AI107052 (MH), P20DK123967 (MH), R01AI168468 (MH), F30AI169748 (SAR) and F30AI150077 (CJB), T32GM007347 (CJB). Confocal laser scanning microscopy in Fig. S1 was performed at the Vanderbilt Cell Imaging Shared Resource (CISR), which is supported by NIH grant DK20593. Some images were created using BioRender.com.

## AUTHOR CONTRIBUTIONS

JRB conceived the study, performed most experiments, and composed the manuscript. SAR, TAB, MAW acquired and analyzed animal data in a blinded fashion. CJB acquired and analyzed all confocal images in figure S1 (blinded). TLR, TLD and AB aided in the generation of the two-component system library. MH conceived the study and oversaw all aspects of its execution. All authors contributed to the generation, analysis, or interpretation of the data and edited the manuscript.

## MATERIALS AND CORRESPONDENCE

Correspondence and requests for materials should be addressed to Maria Hadjifrangiskou.

